# DeepGEOSearch: LLM-Powered Schemaless Retrieval for Biomedical Data Discovery

**DOI:** 10.64898/2025.12.27.696662

**Authors:** Deepshikha Singh, Shashank Jatav, Shefali Lathwal, Arman Kazmi, Soumya Luthra

**Author notes:** DeepGEOSearch Link: https://datafindability.soulb.io Query Data: https://github.com/soulbio/DeepGEOSearch_Data.

## Abstract

Public biomedical repositories contain extensive, valuable datasets, yet identifying datasets that precisely match specific study requirements remains inefficient. Conventional keyword- and schema-based systems frequently fall short when queries encompass multiple biological and experimental facets. To address this, we developed DeepGEOSearch, an LLM-powered, schema-less retrieval system that interprets dataset text directly rather than relying solely on predefined metadata fields. The system extracts key study attributes, harmonizes terminology across sources, and ranks datasets by contextual alignment with natural-language queries while providing verifiable evidence for each match. In this work, we applied DeepGEOSearch for the Gene Expression Omnibus (GEO). The framework is repository-agnostic and can integrate new metadata sources without manual relabeling. This design enables complex compositional queries that existing tools cannot support. In evaluations against strong baselines using a curated evaluation benchmark comprising a mix of query complexities, DeepGEOSearch achieved more than 90% precision and best recall, with the largest performance gains observed on complex, real-world queries. DeepGEOSearch consistently identifies relevant datasets overlooked by conventional search tools, accelerating dataset discovery and improving reuse of public biomedical data.

## 1 Introduction and Background

Reusing public biomedical datasets is essential for reproducibility and new discovery, aligning with FAIR principles [Sansone et al., 2019] that emphasize data findability as foundational for scientific progress. Public repositories house vast amounts of heterogeneous experimental data contributed by research groups worldwide. Yet efficiently locating datasets that satisfy complex, multi-faceted research queries remains surprisingly difficult, limiting the utility of these invaluable resources.

Significant efforts have aimed to improve biomedical dataset discovery through standardized metadata frameworks. The Data Tag Suite (DATS) [dat] and DataMed [Chen et al., 2018] introduced a schema-first paradigm, requiring metadata to follow predefined templates to enable unified indexing and cross-repository search. While this reduced repository heterogeneity, these systems remain dependent on the completeness and accuracy of deposited metadata, which is frequently incomplete or inconsistently recorded. Tools like MetaSRA [Bernstein et al., 2017] mitigate terminological variability by mapping free-text annotations to controlled ontologies, but still require substantial manual or semi-automated curation and cannot fully address unstructured or missing metadata scattered across abstracts and linked publications.

Manual curation efforts, exemplified by GEOMetaCuration [Li et al., 2018], improve metadata quality by enabling domain experts to correct and enhance records. While valuable, such labor-intensive approaches cannot keep pace with the rapid growth of dataset repositories. Restructured GEO [Chen et al., 2019] applies text-mining to extract key facets including disease studied, treatment, sample characteristics, temporal sampling points, and study design elements from narrative descriptions, revealing information hidden beyond structured metadata fields. This partially mitigates metadata gaps but depends on effective natural language processing and cannot handle the evolving complexity and diversity of user queries.

Despite these advances, metadata quality remains a major barrier to dataset discovery. The bioCADDIE benchmark [Cohen et al., 2017] showed that metadata available at deposition are often insufficient to judge dataset suitability, and large-scale audits confirmed anomalies such as uncontrolled field names, missing values, and inconsistent terminology [Gonçalves and Musen, 2019]. As a result, many critical facets remain absent or underrepresented in structured fields [Gonçalves et al., 2017].

Even if metadata were fully normalized, meaning standardized and mapped to consistent ontology or vocabulary terms, retrieval performance would still be constrained by how users query. Bench researchers and clinicians typically submit natural language queries rather than Boolean or ontology-anchored expressions, i.e., structured search commands using logic operators like AND/OR, distinct from plain natural language. Usability studies of DataMed [Dixit et al., 2018] and the bioCADDIE challenge [Roberts et al., 2017] reported that keyword-centric systems fail when relevant attributes are implicit, variably phrased, or inconsistently recorded, resulting in low recall and missed datasets.

*Consider a concrete example: a cancer biologist seeking RNA-seq datasets of obese human patients with renal cell carcinoma treated with a specific drug and sampled at multiple time points*. Such a query includes multiple critical facets such as organism, disease, patient characteristics, intervention, assay, and study design. Existing schemas may capture organism and assay but rarely include detailed patient-level variables such as obesity or treatment exposures. When these details are present, they usually reside in free-text descriptions, abstracts, or linked publications rather than structured metadata fields. Consequently, many relevant datasets remain undiscoverable through schema-constrained or keyword-based search.

These challenges underscore a fundamental limitation: predefined schemas cannot scale with evolving biomedical research or diverse user intents. A paradigm shift is therefore required, one that interprets unstructured information dynamically at query time rather than enforcing rigid metadata at deposition.

Recent advances in transformer-based architectures and Large Language Models (LLMs) enable this shift toward a schema-less architecture. Rather than relying solely on structured metadata fields, these models can extract, normalize, and ground query-relevant attributes dynamically from unstructured text such as titles, abstracts, sample descriptions, and linked publications. This creates a temporary, query-specific schema that adapts flexibly to user intent while maintaining auditability through evidence grounding (tracing information back to evidence phrases in the source text) [Su et al., 2024, Gao et al., 2023, Lála et al., 2023, Beger and Henneking, 2025].

We present *DeepGEOSearch*, an LLM-powered system designed to improve the discovery of biomedical datasets. Unlike traditional schema-dependent search, DeepGEOSearch works flexibly with data described in natural language. It consists of three main components: i) A fast and lightweight search tool that scans Gene Expression Omnibus (GEO) dataset records along with related publication texts to identify candidate datasets quickly. ii) An LLM-driven extractor that reads natural language text to recognize and standardize study attributes for facets such as disease, tissue type, treatment, and cell type. Crucially, this extractor links each identified detail back to its verifiable evidence phrase in the original text, ensuring transparency. iii) A ranking system that orders datasets based on how well they satisfy the user’s query, using clear, evidence-based justifications to prioritize the most relevant results. This approach requires minimal assumptions about how datasets are structured and effectively handles natural language queries blending clinical, experimental, and temporal information. DeepGEOSearch uncovers relevant datasets often missed by traditional schema-dependent searches, unlocking the full potential of public biomedical repositories and making discovery more intuitive and aligned with users’ needs.

We also introduce a general, method-agnostic evaluation benchmark for dataset search systems. This framework uses a balanced set of Short, Medium, and Complex natural language queries to assess search performance comprehensively. It incorporates standard ranking metrics such as normalized Discounted Cumulative Gain (nDCG), Precision, and Recall, alongside user-focused measures such as Time-to-First-Relevant (TTFR) dataset. Crucially, it includes facet-level satisfaction scoring, which evaluates how well the system successfully retrieves datasets by matching attributes to the corresponding query facets (e.g., disease, tissue, cell type). By enabling direct, fair comparisons across different search schemas and models, this evaluation framework guides the development and benchmarking of dataset search tools. We apply this framework exclusively to GEO in our study, consistent with the focus of DeepGEOSearch.

## 2 Methods

### 2.1 Data Findability Workflow

DeepGEOSearch employed a two-stage retrieval pipeline aimed at enhancing the discoverability of biomedical datasets in GEO. See Figure 1.

**Figure 1.**
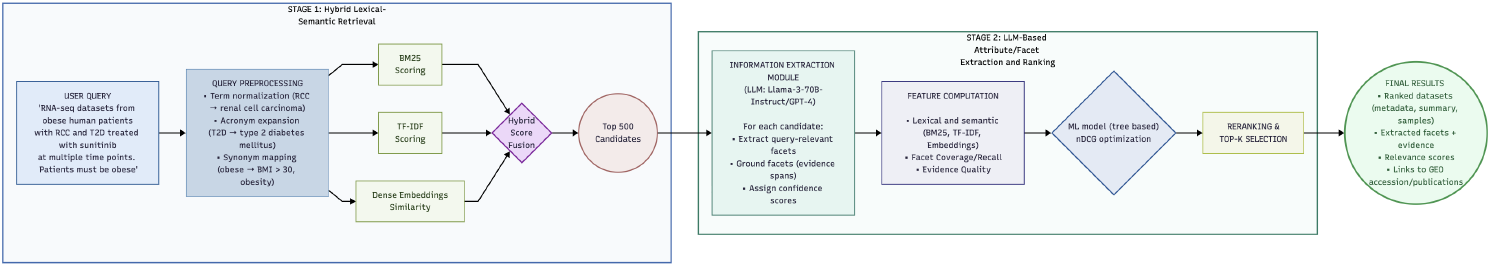
Data Findability Workflow.

#### 2.1.1 Stage 1: Hybrid Lexical-Semantic Retrieval

The first stage combined traditional keyword-based search with semantic understanding. The system scanned dataset titles, summaries, and sample descriptions using a hybrid approach that integrated lexical search techniques (BM25, TF-IDF) with semantic features derived from text embeddings. Lightweight query expansion normalized biomedical terms and common synonyms (e.g., “RCC” → “renal cell carcinoma,” “obese” → “obesity, BMI > 30”) to improve recall. This blended approach captured exact lexical matches important for technical biomedical terms while also accounting for semantically similar expressions, producing a comprehensive shortlist. The output was the top 500 candidate datasets that broadly matched the user’s query both lexically and semantically.

#### 2.1.2 Stage 2: LLM-Driven Attribute/Facet Extraction and Ranking

In the second stage, a large language model processed each candidate to extract and standardize key study attributes for facets such as disease, tissue, treatment, cell type, and patient characteristics. Crucially, every extracted attribute was linked back to specific evidence phrases in the original dataset text (e.g., title, abstract, or sample description) to provide verifiable evidence. This traceability supports user trust and error correction.

##### Key Design Principles

- **Query-Conditioned Extraction:** The model extracted only attributes for the facets relevant to the user query, avoiding unnecessary processing of all metadata dimensions. For example, if the query mentioned *obesity*, the extractor focused on patient characteristics; if it specified *time series*, temporal study-design facets were prioritized.
- **Evidence Grounding:** Every extracted attribute was accompanied by an evidence phrase in the source text (title, abstract, or sample description). Extractions without supporting evidence phrases were automatically discarded to maintain precision and interpretability. This design enabled auditable retrieval, allowing users to verify why a dataset was retrieved and challenge incorrect extractions.
- **Schema-Agnostic Operation:** The extractor did not assume a fixed metadata schema; instead, it dynamically identified context-specific attributes (e.g., disease, organism, assay, cell type, treatment, timepoint, cohort characteristics) based on query semantics. This flexibility surfaced datasets with relevant information in free text, even when structured fields were incomplete.
- **Ranking and Scoring**

The final ranking was computed using a learned scoring function *f* (*x*) defined over grouped features:

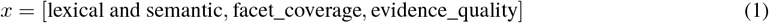

The system reranked candidates by integrating multiple signals: the initial lexical-semantic retrieval scores from Stage 1, how well the extracted attributes met the query requirements, and the confidence scores associated with each attribute’s evidence phrase. A machine-learned ranking model balanced these factors to optimize precision and relevance. This approach ensured that the final results not only matched keyword-level intent but also aligned semantically and contextually with complex research questions.

### 2.2 Query Design and Stratification

We developed a rigorously designed query benchmark to thoroughly evaluate retrieval performance across the varied information needs typical of biomedical dataset searches. This benchmark targeted three key challenges: (1) effective lexical retrieval from sparse text inputs, where minimal query terms must retrieve relevant datasets despite vocabulary variation; (2) satisfying multiple concurrent constraints such as assay type, organism, disease, and experimental design that reflect realistic, complex search demands; and (3) supporting cross-document reasoning when essential metadata like clinical staging, treatment protocols, or outcomes exist primarily in linked publications rather than in structured GEO records. To capture authentic researcher search behavior, all queries were expressed solely in natural language without Boolean operators or special syntax, and each was assigned a difficulty level based on query length, facet complexity, and the extent of cross-document reasoning required.

#### Difficulty Bands

**Short (Lexically Sparse):** Concise queries (2–6 tokens) containing one or two facets (for example, a disease and an assay). These tested lexical recall and synonym and abbreviation handling from minimal textual input.

**Medium (Multi-Faceted):** Mid-length queries (8–20 tokens) encoding an explicit task intent (e.g., “compare cases vs. controls”), alongside multiple constraints such as assay, organism, and optionally a tissue or cell context. These represented the majority of real-world GEO searches, where several constraints must be satisfied simultaneously.

**Complex (Long or Clinical):** Long, detailed queries (20–60 tokens) specifying multiple clinical qualifiers such as stage, severity, timing and, experimental conditions. Because many of these details are often documented only in associated publications, complex queries required reasoning across metadata and full-text evidence, making them particularly suitable for evaluating evidence-aware ranking and evidence grounding.

##### 2.2.1 Mapping Difficulty to Evidence Requirements

Each difficulty band corresponds to distinct evidence sources and retrieval challenges:

**Short** : lexical / surface signals (keywords, synonyms)

**Medium** : contextual signals (co-occurrence across fields)

**Complex** : semantic & temporal signals (linked constraints across sources)

##### 2.2.2 Thematic Coverage and Query Distribution

To ensure broad representativeness, we selected seed topics spanning across major disease areas and modalities represented in the corpus, including breast cancer, lung cancer, COVID-19, Parkinson’s disease, Alzheimer’s disease, type-2 diabetes, and renal cell carcinoma-hypoxia. Common experimental contexts, including bulk RNA-seq, single-cell RNA-seq of peripheral blood mononuclear cells (PBMCs), and case–control study designs were systematically incorporated. For each seed topic, we generated one query per difficulty band (short, medium, complex), yielding a balanced benchmark. This stratification enabled controlled evaluation of retrieval performance as query complexity and reasoning demands increase.

##### 2.2.3 Query Construction Protocol

Query construction followed a structured, multi-step framework:

- Facet Grid Construction: For each topic, relevant facets were enumerated, including disease or condition, assay or modality, organism (preferably human), tissue or cell type, comparison type (e.g., disease vs. control), and clinical qualifiers such as severity, treatment, or timepoint.
- Tiered Query Formulation: Each research intent was rendered in three progressively detailed forms:
  – Short: disease ± assay or cell context.
  – Medium: explicit comparison plus one contextual facet (organism or tissue).
  – Complex: clinical and experimental constraints expressed in natural phrasing rather than Boolean logic.

- Terminology Normalization: Canonical biomedical terms were preferred, but common synonyms and abbreviations were retained to emulate authentic user phrasing (e.g., PBMC - peripheral blood mononuclear cells; COVID-19 - SARS-CoV-2).
- Feasibility Validation: Each query was verified to have at least three plausible relevant datasets in the GEO and linked literature corpus. For complex queries, some facets were confirmed to occur only in abstracts or publications to ensure the need for cross-document reasoning.
- Authorship and Quality Control: Two biomedical researchers independently drafted and reviewed all queries to ensure clarity, facet completeness, and consistent difficulty labeling. Discrepancies were resolved through discussion and consensus.
- Diversity and Non-Redundancy: Near-duplicate queries were excluded. Disease areas and modalities were balanced to avoid domain bias and overrepresentation of specific research subfields.

### 2.3 Ground Truth Preparation

We constructed the relevance ground truth using a pooling-and-triage protocol adapted from TREC evaluation methodology and Microsoft ISE’s efficient ground truth generation framework [Faustino, 2025]. This approach combines multi-system pooling, LLM-assisted triage, and expert annotation to minimize labeling effort while maintaining judgment quality.

We assembled a mixed benchmark comprising (i) public bioCADDIE-style topics and (ii) modern, multi-facet biomedical queries authored by domain experts. These queries captured facets such as assay, disease, biospecimen, and study design. Each query was first expressed in natural language and, when appropriate, paired with an *expert* formulation (Boolean or fielded) to enable comparative evaluation.

#### Pooling and Candidate Selection

For each query, we executed three retrieval systems—DeepGEOSearch, OpenAI Deep Research, and GEO native search. We pooled the *top-k* results from each system (*k* fixed across systems) to construct a candidate set enriched for potentially relevant items while maintaining cross-system fairness.

#### LLM-Assisted Triage

To reduce annotation volume without introducing bias, a general-purpose LLM generated provisional relevance scores with evidence-based justifications for each candidate. Crucially, these scores were used solely to prioritize review order—not as final labels—guiding experts towards likely positives while preserving human judgment as authoritative.

#### Expert Annotation Protocol

Subject-matter experts (SMEs) independently labeled candidates using a structured interface with three relevance levels (Not Relevant, Partially Relevant, Fully Relevant) and facet-satisfaction checkboxes (assay, biosource, condition, comparator, organism). Reviewers had access to (i) repository metadata (GEO series summary, platform, samples), (ii) extracted free-text descriptions, and (iii) linked literature phrases when available. They could also consult full records and PDFs. Disagreements were resolved by a third SME through consensus adjudication.

#### Bias Mitigation

To minimize system-recognition and LLM-induced biases, we implemented three safeguards: (i) blinded system identity during annotation, (ii) randomized candidate order within triage priority bands, (iii) capped per-system contributions to the initial pool per query. SMEs were not shown provisional LLM scores or justifications; these signals served exclusively for queue ordering.

#### Data Release

Final labels were converted to binary and graded relevance formats compatible with metrics such as Precision@k, nDCG, and Recall@k. We publicly release the query set and pooled candidate lists (document IDs only) to support reproducibility and future benchmarking.

## 3 Results

We assessed DeepGEOSearch using the evaluation benchmarks described in subsection 2.2. Performance was measured against two baselines: the native GEO Advanced Search and OpenAI Deep Research. We report Precision@k, Recall@k, nDCG@10, F1, and Time-to-First-Relevant (TTFR), with stratifications by query difficulty and evidence source. Table 1 summarizes system’s overall performance across all queries.

**Table 1.** Benchmark metrics across systems. TTFR = Time-to-First-Relevant.

| System | Number of Datasets Retrieved | TTFR ↓ | nDCG@10 | Precision |  |  | Recall |  |  |
| --- | --- | --- | --- | --- | --- | --- | --- | --- | --- |
|  |  |  |  | P@3 | P@10 | P@20 | R@3 | R@10 | R@20 |
| DeepGEOSearch | 309 | **1.06** | **0.90** | **0.9** | **0.89** | **0.88** | **0.09** | **0.30** | **0.58** |
| GEO | 22 | 4.60 | 0.34 | 0.24 | 0.28 | 0.29 | 0.02 | 0.07 | 0.15 |
| OpenAI | 19 | 1.35 | 0.74 | 0.69 | 0.59 | 0.51 | 0.07 | 0.19 | 0.29 |

Averaged across the benchmarks, DeepGEOSearch achieved P@20 = 0.88, R@20 = 0.58, F1 = 0.70, compared to OpenAI (F1 = 0.37) and GEO Advanced Search (F1 = 0.20) (see Table 1). This corresponds to **89% higher F1 than OpenAI** and **350% F1 over GEO**. DeepGEOSearch substantially outperformed both baselines.

Table 2 stratifies results by difficulty band, revealing consistent DeepGEOSearch advantages that amplify with query complexity.

**Table 2.** Performance by query category and systems. TTFR = Time-to-First-Relevant.

| Category | System | Number of Datasets Retrieved | TTFR ↓ | nDCG@10 | Precision |  |  | Recall |  |  |
| --- | --- | --- | --- | --- | --- | --- | --- | --- | --- | --- |
|  |  |  |  |  | P@3 | P@10 | P@20 | R@3 | R@10 | R@20 |
| short | DeepGEOSearch | 83 | **1.0** | **0.94** | **0.89** | **0.93** | **0.92** | **0.1** | **0.33** | **0.65** |
|  | GEO | 20 | 6.7 | 0.28 | 0.11 | 0.10 | 0.10 | 0.01 | 0.03 | 0.07 |
|  | OpenAI | 19 | **1.0** | 0.64 | 0.56 | 0.57 | 0.59 | 0.06 | 0.20 | 0.39 |
| medium | DeepGEOSearch | 500 | **1.1** | **0.90** | **0.93** | **0.89** | **0.88** | **0.07** | **0.22** | **0.42** |
|  | GEO | 23 | 2.0 | 0.47 | 0.41 | 0.44 | 0.46 | 0.03 | 0.11 | 0.22 |
|  | OpenAI | 18 | 1.3 | 0.76 | 0.74 | 0.69 | 0.63 | 0.05 | 0.17 | 0.28 |
| complex | DeepGEOSearch | 100 | **1.0** | **0.88** | **0.87** | **0.86** | **0.85** | **0.13** | **0.42** | **0.83** |
|  | GEO | 20 | 10.3 | 0.14 | 0.00 | 0.10 | 0.10 | 0.00 | 0.04 | 0.08 |
|  | OpenAI | 20 | 1.6 | 0.76 | 0.67 | 0.44 | 0.26 | 0.1 | 0.21 | 0.26 |

### 3.1 Recall by Query Complexity

#### Short (lexically sparse)

DeepGEOSearch maintained high precision with competitive recall (P@20 = 0.92, R@20 = 0.65, F1@20 = 0.76), outperforming OpenAI (F1@20 = 0.47, + 62%) and GEO advanced search (F1@20 = 0.08, + 950%).

#### Medium (multi-faceted)

Although recall declined across all systems, reflecting increased constraint specificity, DeepGEOSearch retained the strongest overall performance (P@20 = 0.88, R@20 = 0.42, F1@20 = 0.57), surpassing OpenAI (F1@20 = 0.38, + 50%) and GEO advanced search (F1@20 = 0.30, + 90%).

#### Complex (long or clinical)

DeepGEOSearch demonstrated its largest performance margin (P@20 = 0.85, R@20 = 0.83, F1@20 = 0.84), exceeding OpenAI (F1@20 = 0.26, + 223%) and GEO advanced search (F1@20 = 0.09, + 900%).These gains stem primarily from substantially higher recall on literature-anchored intents while maintaining precision ≥ 0.85, confirming the value of evidence-aware ranking and cross-document reasoning for clinically constrained queries.

Across short to complex queries, DeepGEOSearch’s precision remained consistently high (0.85–0.92) while recall varied with intent specificity (0.65 short, 0.42 medium, 0.83 complex). Recall declined in medium category, where DeepGEOSearch retrieved a large number of relevant datasets, though not all present in the Top-20, the majority were present within the Top-100. OpenAI’s recall remained low (0.26) with reduced precision on complex queries, and GEO underperformed on both precision and recall across all categories. Taken together, DeepGEOSearch consistently outperformed the baselines (Table 2). OpenAI’s semantic retrieval surpassed GEO on every split (overall F1 = 0.37 vs 0.20) but remained limited on complex, text-heavy intents, precisely where the DeepGEOSearch’s PubMed/PMC linking and evidence-aware ranking contribute the largest gains. For users issuing complex, clinically framed queries, DeepGEOSearch retrieved substantially more relevant datasets without loss of precision, thereby reducing downstream triage effort. For short queries, high precision paired with strong recall indicates robust handling of sparse evidence signals.

### 3.2 Ranking Quality and Early Precision/Recall

DeepGEOSearch achieved the highest early-ranking quality (Table 2). Average nDCG@10 was 0.94 for short, 0.90 for medium, and 0.88 for complex queries, compared to OpenAI’s 0.64 / 0.76 / 0.76 and GEO’s 0.28 / 0.47 / 0.14. These results confirm that DeepGEOSearch consistently surfaces the strongest supporting evidence within the Top-10, particularly for complex, prose-anchored intents derived from literature sources.

Per-query trends (nDCG@10 vs. *pooled* R@20 - fraction of all relevant datasets across systems recovered at depth K) followed a consistent pattern (see Table 2):

- DeepGEOSearch points clustered at high nDCG@10 (short/medium/complex: **0.94**/**0.90**/**0.88**) with competitive pooled recall at *K*=20 (0.65/0.42/0.83), indicating concentrated early relevance that continues to accrue with depth.
- OpenAI exhibited lower nDCG@10 (0.64/0.76/0.76) and correspondingly lower pooled R@20 (0.39/0.28/0.26), suggesting that more of its relevant items reside deeper but remain fewer overall.
- GEO showed both lower nDCG@10 (0.28/0.47/0.14) and the weakest pooled recall (0.07/0.22/0.08).

### 3.3 Precision@K (Relevance Density)

Across *K* = 1–20, DeepGEOSearch maintained high precision whereas both baselines degraded sharply (Figure 2).

**Figure 2.**
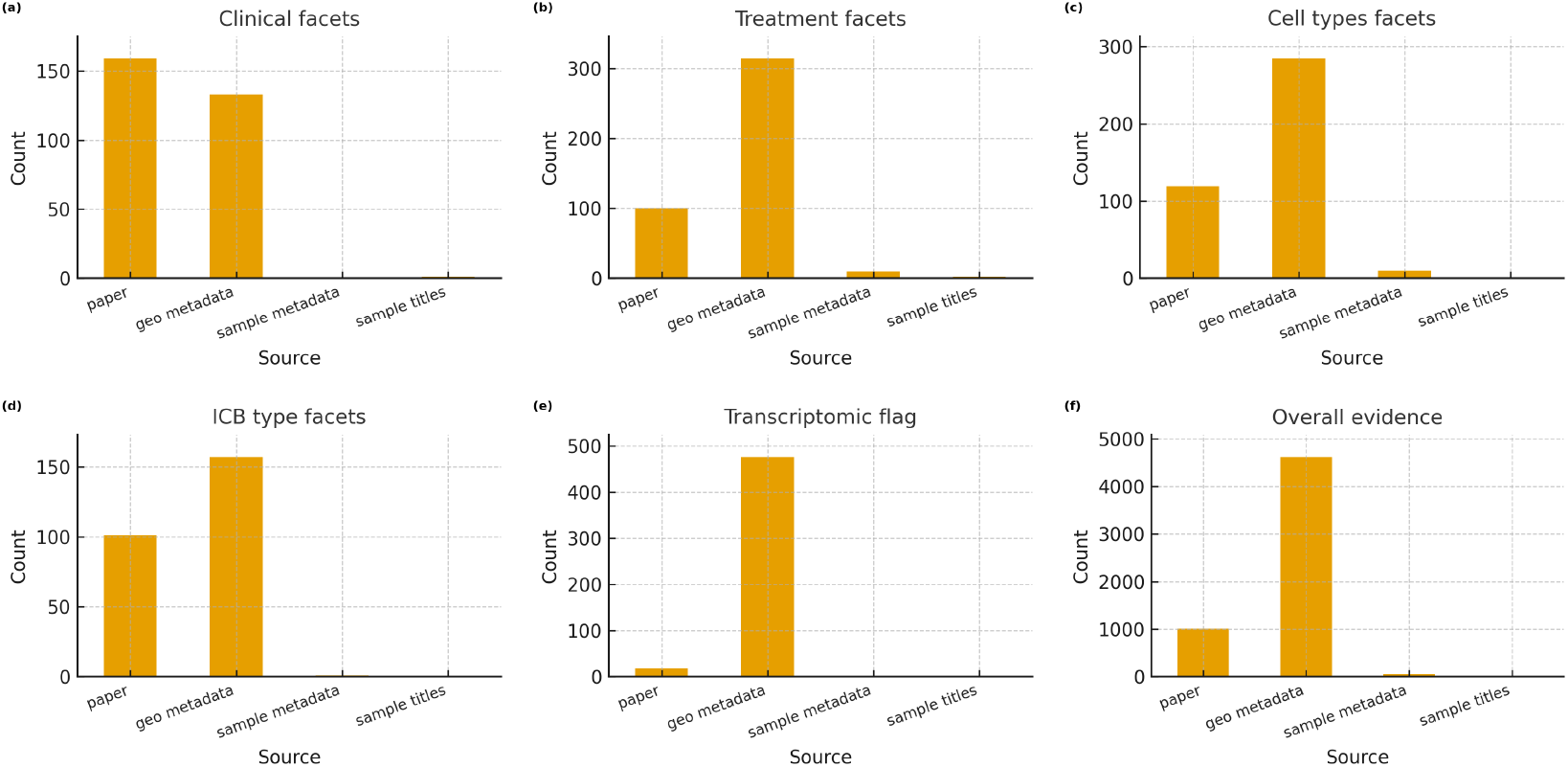
Distribution of evidence sources across facets. Panels (a–e) show aggregated evidence counts for a representative medium query (*Datasets profiling transcriptomic responses to immune checkpoint blockade in melanoma*), stratified by the following facets: clinical stage/severity, treatment regimen, cell type, immune checkpoint blockade (ICB) type, and transcriptomic flag. Panel (f) shows the aggregate distribution across all benchmark queries. Evidence sources include GEO metadata (mentioned on GEO dataset summary page), sample metadata (metadata for each sample on GEO), sample titles, and linked literature (paper).

- **Complex:** DeepGEOSearch ≈ 0.86 plateau (vs. OpenAI 0.67 *→* 0.26, GEO 0.00 *→* 0.10).
- **Medium:** DeepGEOSearch ≈ 0.88–0.93 (vs. OpenAI 0.74 *→* 0.63, GEO 0.41 *→* 0.46).
- **Short:** DeepGEOSearch ≈ 0.9 plateau (vs. OpenAI 0.56 *→* 0.59, GEO 0.1 *→* 0.11).

This demonstrates that DeepGEOSearch preserves relevance density deeper into the ranking, reducing triage burden for users.

### 3.4 Recall@K (Coverage)

We report pooled recall. Recall curves increase with *K* but do not saturate at 1.0 by *K*=20; instead they differentiate systems and difficulty bands (Table 2).

- **Complex:** DeepGEOSearch achieved the highest pooled recall at depth (R@20 = **0.83**), with early coverage R@3 = **0.13** and R@10 = **0.42**. OpenAI trailed (0.10, 0.21, 0.26), and GEO remained substantially lower (0.00, 0.04, 0.08). This depth advantage aligns with DeepGEOSearch’s superior precision (P@3/10/20 = **0.87**/**0.86**/**0.85**) and nDCG@10 = **0.88**.
- **Medium:** DeepGEOSearch led across depths (R@3/10/20 = **0.07**/**0.22**/**0.42**), followed by OpenAI (0.05/0.17/0.28) and GEO (0.03/0.11/0.22). Precision remained highest for DeepGEOSearch (0.93/0.89/0.88) with the best nDCG@10 = **0.90**.
- **Short:** DeepGEOSearch achieved stronger coverage (R@3/10/20 = **0.10**/**0.33**/**0.65**) compared to OpenAI (0.06/0.20/0.39) and GEO (0.01/0.03/0.07). Its precision remained high (0.89/0.93/0.92), with highest nDCG@10 at **0.94**.

Taken together, DeepGEOSearch continues to uncover relevant datasets even deeper in the ranking while maintaining the highest early precision and nDCG across difficulty bands.

### 3.5 Time-to-First-Relevant (TTFR)

TTFR (lower is better) was best or tied-best for DeepGEOSearch across all query bands (short: **1.0**; medium: **1.1**; complex: **1.0**), matching or improving upon OpenAI and substantially outperforming GEO. This demonstrates that evidence-aware retrieval does not compromise latency.

## 4 Analysis

To understand the mechanisms driving the performance gains reported in section 3, we conducted a qualitative audit of the retrieval process. This analysis focuses on the distribution of retrieval signals across sources and the system’s capability to ground retrieval in specific evidence phrases.

### 4.1 Impact of Evidence Sources

We analyzed the origin of the evidence phrases used by DeepGEOSearch to justify relevance. Evidence was classified into two primary sources:

1. **Repository Metadata:** Structured fields such as title, summary, and sample characteristics.
2. **Linked Literature:** Unstructured full-text or abstracts from PubMed/PMC articles associated with the dataset.

#### Distribution of Information

Information is not uniformly distributed across these sources. As shown in Figure 2(a–f), technical facets—such as Assay Type, Organism, or Cell Line—are reliably present in Repository Metadata. However, clinical and outcome-related facets are frequently absent from metadata fields. For example, when analyzing datasets related to Immune Checkpoint Blockade (ICB), we observed that clinical variables like *Treatment Response* (e.g., “Non-responders”) or Disease Stage (e.g., “Stage IV”) were recorded both in GEO metadata and linked literature as opposed to technical attributes like *transcriptomic flag* that are present exclusively in GEO metadata.

#### Explaining the Performance Gap

This divergence explains why DeepGEOSearch outperforms baselines on complex queries:

- **Metadata-Centric Baselines:** Systems indexing only metadata (such as GEO Advanced Search) fail on clinical queries because the relevant signals are absent from the indexed fields.
- **DeepGEOSearch:** By dynamically ingesting the Linked Literature, the system recovers this “invisible” evidence, effectively performing cross-document reasoning to connect a dataset ID to the clinical narrative in the associated publication.

### 4.2 Case Studies in Evidence Grounding

A key architectural feature of DeepGEOSearch is Evidence Grounding, the requirement to verify a match by citing the exact evidence phrase that satisfies a query constraint. This ensures transparency and prevents hallucination. The following examples illustrate this capability.

#### Case 1: Resolving Implicit Clinical Outcomes

- **Query:** “*Biomarkers for immunotherapy response in NSCLC*.” Figure 3
- **Challenge:** Keyword search retrieves generic immunotherapy datasets regardless of whether the study measured patient response.
- **DeepGEOSearch Resolution:** The system successfully identified relevant datasets by locating explicit evidence of outcome measurement. For instance, it extracted the phrase *“patients were stratified into responders and non-responders according to RECIST 1*.*1”* from the Methods section of the linked literature.

**Figure 3.**
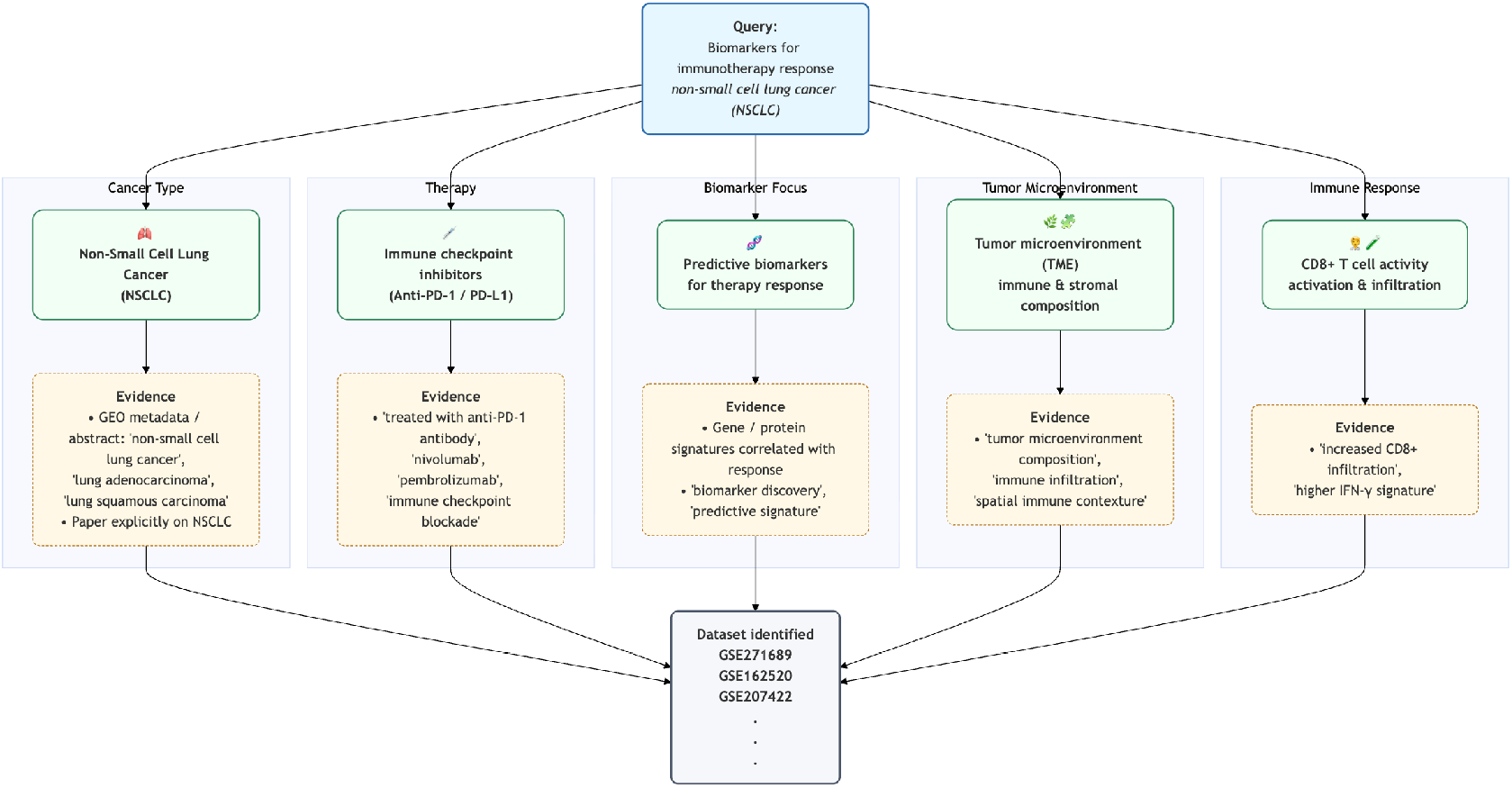
Evidence grounding for a representative medium query: *Biomarkers for immunotherapy response in NSCLC*. The schematic illustrates how datasets are identified by verifying evidence across five key facets: cancer type, therapy, biomarker focus, tumor microenvironment (TME), and immune response. Representative evidence phrases extracted from metadata confirm specific query attributes to validate dataset relevance.

By grounding the retrieval in this specific evidence phrase, the system filtered out non-specific studies and prioritized those with usable phenotypic data.

#### Case 2: Locating Granular Variables

- **Query:** “*Type 2 diabetes; adipose tissue; HbA1c reported*.”
- **Challenge:** “HbA1c” is a clinical variable rarely indexed in GEO sample tags.
- **DeepGEOSearch Resolution:** The system retrieved datasets where the linked literature’s Patient Characteristics Table explicitly reported HbA1c values (e.g., “*Mean HbA1c: 8*.*2%*”). The system highlighted this evidence phrase to justify relevance. This confirms that for clinical queries, the effective “schema” is often constructed on-the-fly from the literature tables and text, as the necessary metadata fields simply do not exist in the repository.

## 5 Discussion

The rapid expansion of biomedical data has exposed a fundamental scalability constraint in conventional discovery systems: scientific inquiry evolves more quickly than repository standards can adapt. Schema-dependent retrieval methods rely on fixed metadata fields that inevitably lag behind emerging experimental modalities and clinical terminology. Our results indicate that DeepGEOSearch mitigates this mismatch by operationalizing a schema-less retrieval paradigm. By shifting the burden of structural interpretation from the data depositor to the retrieval algorithm, the system adapts dynamically to user intent rather than forcing novel research concepts into static, legacy schemas.

This architectural shift directly addresses the semantic gap highlighted by our analysis. Whereas technical attributes (such as organism or assay) are consistently indexed, high-value clinical context, such as disease stage, treatment response, and longitudinal outcomes, frequently resides only within the unstructured prose of linked literature. DeepGEOSearch bridges this gap through a hybrid design that couples the efficiency of keyword search with LLM-driven semantic reasoning and evidence grounding. This dual strategy explains the system’s substantial performance advantage on complex, literature-grounded queries (F1 = 0.84 vs 0.26), demonstrating that effective biomedical retrieval requires interrogation of full scientific text rather than reliance on metadata alone.

### Limitations

Despite its strengths, the current implementation has inherent limitations. First, although the system emphasizes unstructured text analysis, retrieval quality remains partly dependent on the richness of dataset descriptions and the availability of linked literature; sparse documentation can limit evidence extraction and impact recall. Second, while LLM-driven attribute extraction improves accuracy over keyword matching, it may yield inconsistent confidence estimates when processing highly noisy or ambiguous metadata. Finally, the multi-stage architecture, necessary for deep semantic reasoning, introduces additional computational latency relative to purely lexical systems, which may affect real-time usability for very large candidate sets.

### Future Directions

Addressing these challenges will guide future development. Domain-specific tuning, lightweight re-ranking layers, and caching strategies offer promising avenues to balance semantic precision with system responsiveness. More broadly, the schema-less design provides inherent adaptability: because schemas are constructed on demand, the system can accommodate emerging disease subtypes or therapeutic classes without costly re-curation. This flexibility positions the framework as a generalizable interface for cross-repository search, enabling integration across heterogeneous resources (e.g., ArrayExpress, ENA) and advancing the practical realization of FAIR principles.

## 6 Conclusion

We demonstrate that the major barriers to biomedical dataset discovery arise not from the data itself but from the constraints of traditional, schema-dependent retrieval systems. DeepGEOSearch overcomes these limitations through a schema-less, LLM-powered framework that connects the rigid structure of repositories like GEO with the complex natural language of scientific inquiry. By combining semantic reasoning with fast lexical search, it exposes clinical context such as outcomes and longitudinal details that would otherwise remain hidden in linked literature. Our results suggest that future discovery tools must move beyond static indexing toward adaptive systems that align with how scientists reason. DeepGEOSearch validates this paradigm, ensuring that datasets are prioritized by the scientific value of their content rather than the completeness of their metadata.

